# *Chlamydia* iron starvation links nutritional immunity to pathogen recognition

**DOI:** 10.1101/2025.08.08.669282

**Authors:** Monisha R. Alla, Nick D. Pokorzynski, Junghoon Lee, Scot P. Ouellette, Rey A. Carabeo

## Abstract

Nutritional immunity is an antimicrobial strategy that evolved to starve pathogens of essential nutrients, with death as the desired outcome. Here, we report that transient iron starvation of the obligate intracellular pathogen *Chlamydia trachomatis*, growing in endocervical epithelial cells, enhances pathogen recognition by the host cell through the dysregulation of a peptidoglycan (PG) remodeling enzyme, resulting in the activation of the nucleotide-binding oligomerization domain 2 (NOD2) pathway that recognizes PG fragments, increased production of tumor necrosis factor alpha (TNF) via increased activation of NF-κB, which correlated with death of infected cells. Activation of the NOD2/ NF-κB signaling axis is linked to the dysregulated overexpression of the PG remodeling enzyme AmiA and the subsequent cleavage and mislocalization of D-Ala-D-Ala analog. Inhibiting *amiA* transcriptional upregulation by CRISPR interference reduced pathogen recognition. We propose that nutritional immunity in general mediate abnormal expression of bacterial genes linked to pathogen-associated molecular patterns.

**Importance:** Limiting pathogen access to essential nutrients is the central tenet of nutritional immunity, with the outcome being severe starvation and eventual death of the pathogen. However, pathogen starvation induces several physiological changes prior to its death. They include errors in several biological processes, including metabolism and gene expression, which could lead to pathogen death. Here, we demonstrate that iron starvation of the clinically relevant human pathogen *Chlamydia trachomatis* significantly dysregulates the expression of a peptidoglycan remodeling amidase, AmiA to enhance chlamydial recognition by the host cell and the subsequent increased production of tumor necrosis factor and death of infected cells to the detriment of *Chlamydia*.

## Introduction

The concept of nutritional immunity involves killing of the pathogen via nutrient withdrawal, which the host achieves through a variety of mechanisms (1). They include upregulation of catabolic enzymes (2, 3), nutrient-sequestering factors (4-6), down-regulation of expression of acquisition systems, etc. The net result is the starvation of the pathogen followed by death. At the same time, the pathogen through evolution has devised strategies to counteract this. For example, iron sequestration by host iron-binding proteins is overcome by the synthesis of high-affinity siderophores and the corresponding upregulation of the iron-siderophore complex retrieval system (7, 8). These strategies are induced during infection, which represents a significant environmental change for the pathogen. The obligate intracellular bacterium *Chlamydia trachomatis* is a clinically relevant pathogen of humans (9). Because of its relatively small genome (1 Mbp), which through reductive evolution has led to the loss of multiple metabolic genes among others, C. trachomatis relies on the host for the majority of its nutritional requirements (10-12). This dependence represents a major vulnerability of C. trachomatis in that host-mediated nutrient limitation (*i*.*e*., nutritional immunity) is very effective in limiting growth and replication of the bacteria. For example, chlamydial growth inhibition by interferon-gamma (IFNγ) is associated with significant reduction in the levels of tryptophan via degradation by the IFNγ-induced enzyme indoleamine 2,3-deoxygenase (IDO1) (13-16). Tryptophan catabolism appears to be the major anti-chlamydial effector of IFNγ in that growth and replication of the pathogen is largely unaffected during IFNγ exposure when IDO1 activity is absent, either through inhibition by levo-1-methyltryptophan (L-1MT) or genetic ablation (17). This is despite the presence of other proposed IFNγ effectors. An outcome of tryptophan starvation is the induction of an abnormal growth state, termed persistence, which is marked by aberrant morphology, the absence of cell division, and a genome wide transcriptional dysregulation with concomitant discordant translation (18-20). Persistent chlamydiae are developmentally arrested, with the crucial transition of the vegetative, but non-infectious reticulate bodies (RBs) to the metabolically quiescent infectious elementary bodies (EBs) inhibited. In other words, persistent chlamydiae remain non-infectious. Starving *C. trachomatis* of other nutrients, including iron also leads to a similar persistent state marked by the same morphological and transcriptomic changes (21). The pathogenic relevance of persistent chlamydiae is unclear, but they have been observed in clinical samples from infected individuals (22). *In vitro*, persistence is reversible, with supplementation of nutrients or withdrawal of growth inhibitory cytokines restoring chlamydial growth and development (23). Reactivation of persistent chlamydiae *in vivo* has not been reported. The ability of *C. trachomatis* to achieve a viable but non-replicating growth state both *in vitro* and *in vivo* indicate that death of the pathogen is not necessarily the outcome nutritional starvation or this bacterium. To gain a more in-depth insight into chlamydial stress adaptation and its implications on its relationship with its host epithelial cell, data from dual RNA sequencing experiments to define the chlamydial and host cell transcriptomes during transient (16h) or sustained (24h) starvation of either iron or tryptophan led to a novel interpretation of chlamydial persistence. Our findings indicate that *C. trachomatis* transitions from a stressed to persistent growth state, and this transition is reflected by marked changes to the transcriptomes unique to each stress. These stress-specific features are lost in persistent chlamydiae, where genomewide transcriptomes displayed significant overlap in differentially expressed genes. This is consistent with a corruption in the adaptive transcriptional response to stress. In other words, persistence is likely a symptom of *C. trachomatis* being overwhelmed by either a single nutritional stress sustained for long periods or by multiple stresses induced simultaneously. This conclusion contrasts with the long-held notion that persistence is a chlamydial stress adaptation strategy (24). This then raises the significance of the transcriptomic response to transient nutritional starvation. We observed that both stresses induced expression of genes associated with synthesis of metabolic precursors, as well as components of the translation machinery in *C. trachomatis* (21). We previously interpreted this transcriptomic profile as being part of the chlamydial prioritization strategy to promote efficient recovery from iron starvation, and it is possible that the same applies to tryptophan starvation. Of particular interest were the sets of chlamydial and host genes that were differentially expressed only in one stress condition (24), which raises the possibility of certain aspects of pathogen-host cell interaction being unique to a particular stress. These observations are addressed further here to define their biological significance to *Chlamydia*-host interactions. We focused on the transcriptional induction of *TNF1*, which was unique to iron-starved C. trachomatis-infected HeLa monolayers. Infection of HeLa cells with *C. trachomatis* induces TNF production, but its significance has not been fully addressed. Indeed, studies that examine TNF effects on *C. trachomatis*-infected cells often include its addition to the growth media (25-27). This mimics effects of TNF from exogenous sources, instead of TNF produced by the infected epithelial cell.

In this report, we define the molecular basis for TNF production by infected epithelial cells (*e*.*g*. HeLa and the endocervical cell line End1) in the context of iron starvation, a stress that significantly enhances TNF expression and production. Iron starvation dysregulates the expression of a peptidoglycan remodeling enzyme AmiA, an amidase that cleaves the D-Ala-D-Ala motif during *C. trachomatis* cell division. AmiA overexpression is required for the mislocalization of a Click-It chemistry-compatible analog of D-Ala-D-Ala (EDA-DA) that was pre-incorporated in the PG network prior to iron starvation. Inhibition of AmiA overexpression maintained bacterial localization of the EDA-DA and reduced NF-κB nuclear translocation, transcription of *TNF1*, production of TNF, and the incidence of cell death in the infected cell population. Collectively, the data point to an enhanced pathogen recognition by infected epithelial cells due to the dysregulated expression of a PG-processing enzyme that consequently generated PG intermediates recognized by the host cell. A broader implication is the refinement of the nutritional immunity paradigm that includes induction of gene expression errors to expose the pathogen to host recognition machinery.

## Results

### TNF production is unique to iron-starved *C. trachomatis*-infected endocervical cells

From previous dual RNA sequencing experiments and stress survey of infected HeLa cells starved for either iron or tryptophan, TNF1 overexpression was found to be uniquely associated with iron starvation with the iron chelator 2,2-bipyridyl (Bpdl) (Supplemental Fig. 1A (24). To confirm this finding, the endocervical epithelial cell line E6/E7-immortalized End1 was infected with *C. trachomatis* serovar L2 (*CtrL2*) at a multiplicity of infection (MOI) of 5 for 8 h, followed by treatment with Bpdl for 16 h. At 24 h post-infection (hpi), conditioned media and cell lysates were collected for a comprehensive cytokine profiling by Multiplexing LASER Bead Technology (Fig. 1A). Elevated levels of TNF were observed by ELISA, which were confirmed by western blotting of cell lysates and RT-qPCR quantification of *TNF* transcripts (Fig. 1 B & C). Increased expression and production of TNF were observed in Bpdl-treated *CtrL2*-infected cells, but not in infection-alone or Bpdl-only treated samples. In addition, similar results were obtained during infection with the genital serovariant *C. trachomatis* serovar D (*CtrD*) (Supplemental fig. 1B). Because TNF has cytocidal properties, the level of cell death was monitored by fluorescence imaging of samples stained with Annexin V to detect phosphatidylserine (PS) translocation to the outer leaflet of the plasma membrane (28, 29). As observed in lactate dehydrogenase levels found in the conditioned media (Fig. 1D), along with qualitative analysis of representative images of annexin V-stained samples shown in Fig. 1E and Supplemental fig. 2, culture conditions that promoted enhanced TNF production correlated with increased cell death. While increased TNF was not implicated directly in increased incidence of cell death, the observations were consistent with known effects of TNF (30).

**Figure 1.**
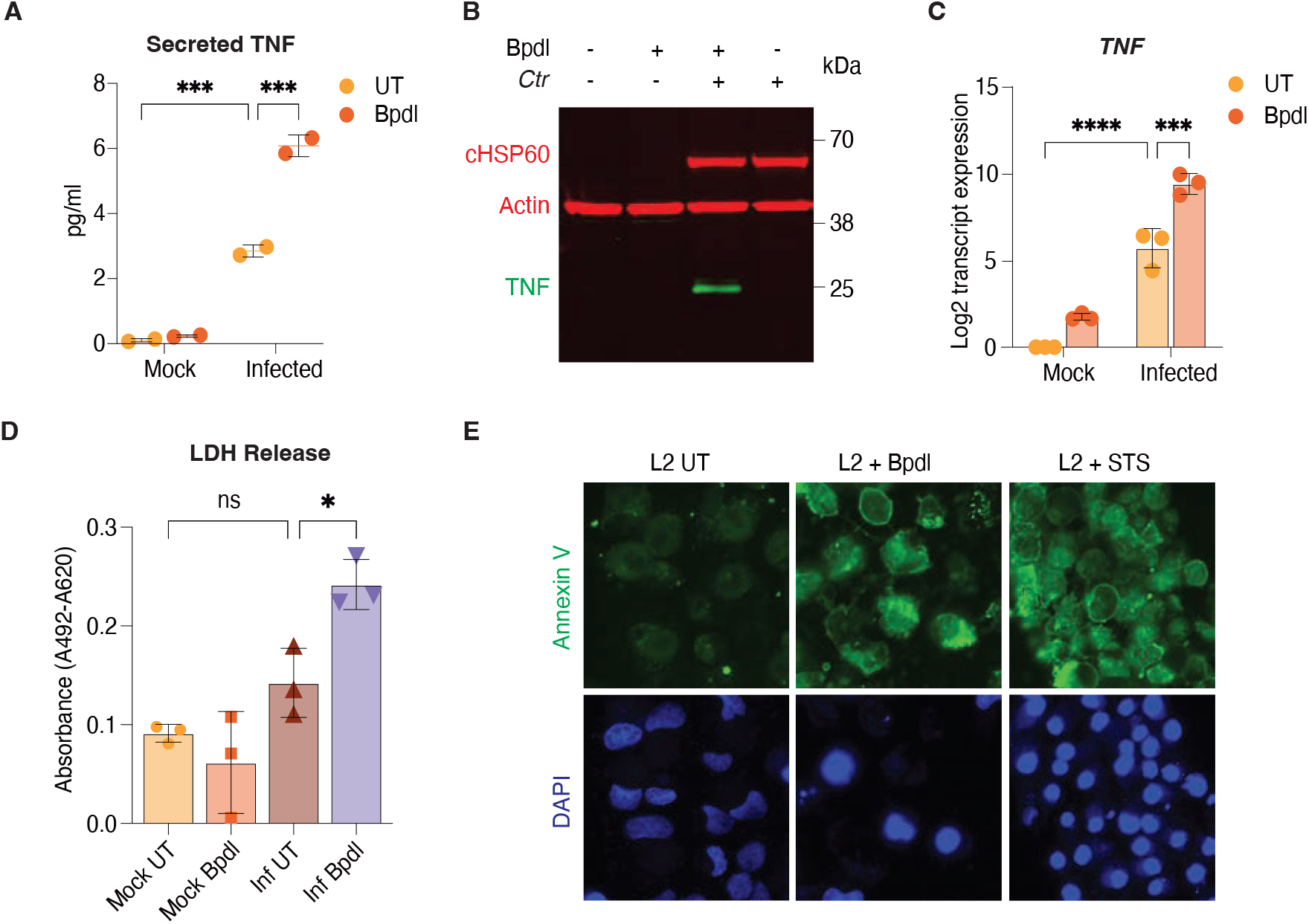
*CtrL2*-infection under iron-starved conditions enhances TNF expression and cell death in epithelial cells. **A)** TNF levels in Hela cell supernatants measured at 24hpi using Multiplexing LASER Bead Technology in mock- or *CtrL2*-infected cells, with or without bipyridyl treatment (100µM), (n=2). **B)** Representative western blot analysis of intracellular TNF in End1 cells at 24hpi under the same conditions. Results were independently confirmed in HeLa cells as well (n=2). **C)** TNF mRNA expression in HeLa cells at 24hpi analyzed by RT-qPCR normalized to HPRT1 (n=3). **D)** LDH release measured in HeLa cell supernatants at 24hpi, shown as absorbance, indicating cytotoxicity under the same infection and treatment conditions (n=3). **E)** Representative micrographs of HeLa cells stained with Annexin V at 24hpi following *CtrL2* infection and bipyridyl treatment (n=2). Statistical significance determined by two-way ANOVA and Tukey’s multiple comparisons. ****p<0.0001; ***p<0.001; ns, not significant.

**Figure 2.**
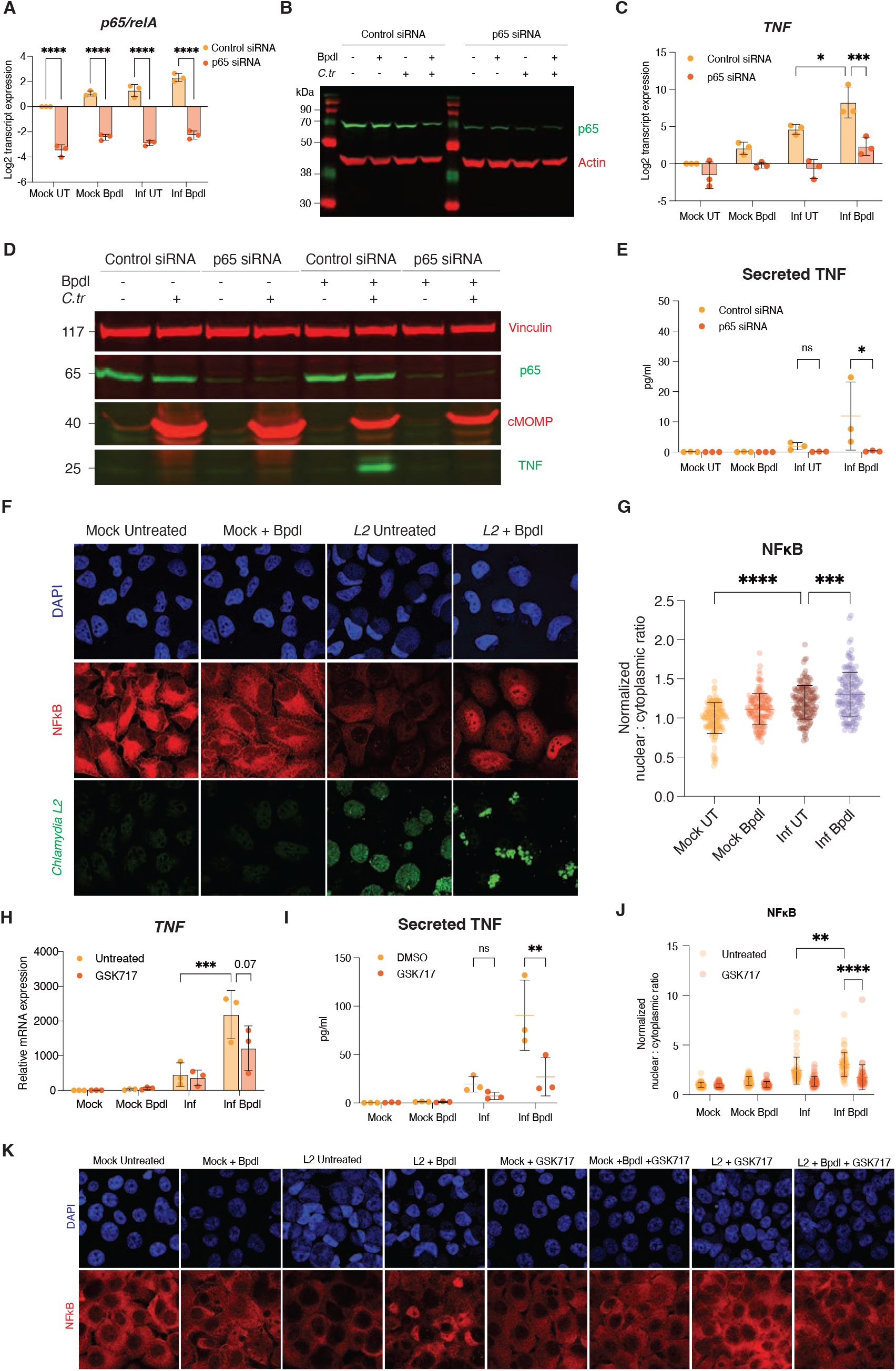
Iron starved *CtrL2* infected epithelial cells induce increased NF_κ_B activity via NOD2 signaling pathway. **A)** p65 mRNA expression in HeLa cells at 48h post-transfection, comparing mock and infected cells, with or without bipyridyl treatment (n=3). **B)** Western blot analysis of intracellular p65 protein levels in HeLa cells at 48h post-transfection using β- actin as loading control (n=2). **C)** *TNF* transcript levels in HeLa cells at 48h post-transfection comparing mock and *CtrL2*-infected cells under untreated or iron-starved conditions, as measured by qPCR (n=3). **D)** Intracellular *TNF* expression in HeLa cells with p65 knockdown at 48h post-transfection detected by western blotting, using vinculin as the loading control and chlamydial MOMP as the infection control (n=1). **E)** TNF protein levels in HeLa cell supernatants at 48h post-transfection measured using Multiplexing LASER Bead Technology in mock-or *CtrL2*-infected with p65 knockdown, with or without bipyridyl treatment (100µM), (n=3). **F)** Micrographs representing NFκB (red) translocation to the nucleus (blue) in uninfected and *CtrL2* infected (green) HeLa cells. **G)** Quantification of NFκB nuclear translocation in (F) expressed as the ratio of nuclear to cytosolic NFκB fluorescence. 50 random cells were measured per treatment group. **H)** *TNF* transcript levels in GSK717-pretreated End1 cells analyzed by qPCR at 24hpi under infected and iron-starved conditions (n=3). **I)** TNF levels in GSK717 pre-treated End1 cell supernatants at 24hpi, measured using Multiplexing LASER Bead Technology in mock-or *CtrL2*-infected cells, with or without bipyridyl treatment (n=3). **J)** Quantification of NFκB nuclear translocation in End1 cells expressed as nuclear to cytoplasmic ratios for the micrographs shown in **K)** when GSK717-pretreated End1 cells were infected under untreated and iron-starved conditions. 50 random cells were measured per treatment group. Statistical significance determined by two-way ANOVA and Tukey’s multiple comparisons test. ****p<0.0001; ***p<0.0002; **p<0.002; *p<0.03; ns, not significant.

### Activation of NF-κ B is required for enhanced TNF production by iron-starved *C. trachomatis*-infected endocervical cells

The promoter region of human *TNF1* gene contains binding sites of several transcription factors, including NF-κB (31-34). To determine if *TNF1* regulation is linked to NF-κB, the p65 subunit was knocked down in expression by transfection of short-interfering RNA (siRNA). At 48 h post-transfection, knockdown was confirmed by RT-qPCR (Fig. 2A) and western blot (Fig. 2B) in the experimental groups. Both Bpdl-treatment and infection of control siRNA-transfected samples separately resulted in approximately a two-fold increase in p65 expression relative to mock-infected/mock-treated control, but the respective differences in levels did not reach statistical significance. The combination of infection and Bpdl treatment increased p65 expression by four-fold. Despite these increases, knockdown was efficient regardless of infection and/or treatment conditions, with p65 transcript levels reduced by at least 90% by 24 h post-transfection. Levels of *TNF1* transcripts (Fig. 2C), and total and secreted TNF (Fig. 2D, 2E) were monitored in the indicated samples transfected with either control or p65 siRNA. As shown in Figs. 2C-E, p65 knockdown decreased *TNF1* and TNF levels, consistent with an important role for NF-κB in this process.

As further confirmation, levels of nuclear NF-κB were quantified by NIH ImageJ from images acquired by immunofluorescence assay and confocal microscopy imaging (Fig. 2F). Creation of masks of nuclei were guided by DAPI images, and fluorescence intensities of at least 10 nuclei and the corresponding cytosolic compartments from three independent samples were measured, with data expressed as fluorescence ratios of nuclei-to-cytosol intensities. In *CtrL2* infection-only samples, there was a statistically significant increase in levels of nuclear NF-κB when compared to mock-infected/mock-treated control (Fig. 2G). Further increase was observed in *CtrL2*-infected and Bpdl-treated samples, relative to infection alone. We conclude that the dependence of TNF expression on NF-κB is linked to nuclear localization (*i*.*e*., activation) of this transcription factor.

### Activation of NF-κB and the subsequent enhanced TNF production by iron-starved *C. trachomatis*-infected endocervical cells is NOD2-dependent

One of the genes upregulated in the infected and Bpdl-treated cells is NOD2. Because NF-κB is activated by NOD2 in the presence of pathogen-associated molecular patterns (PAMPs), we evaluated the role of this pathogen recognition receptor (PRR) in both NF-κB activation via changes in nuclear-to-cytoplasmic intensity ratios and TNF levels in the conditioned media under the conditions specified (Fig. 2H-K) (35, 36). Knockdown of NOD2 was not technically feasible in End1 cells. Therefore, inhibition by the reported NOD2-specific small molecular inhibitor GSK717 was used to determine its role in enhanced *TNF1* transcription, secretion, and NF-κB nuclear translocation (37). As shown in Fig. 2H, GSK717 treatment decreased *TNF1* transcripts relative to the parallel mock-treated control, albeit not reaching a p<0.05 threshold. Despite this marginal result, TNF secretion was significantly reduced by GSK717 treatment (Fig. 2I). In addition, nuclear translocation of NF-κB was significantly reduced (Fig. 2J, 2K). Collectively, the data are consistent with NOD2 playing an important role in activating NF-κB in and enhancing TNF production by iron-starved, *CtrL2*-infected End1 cells.

### *De novo* protein synthesis by iron-starved *C. trachomatis* is required for NF-κB activation and enhanced TNF production

We next examined the role of *CtrL2* in NF-κB activation and enhanced TNF production, specifically the potential requirement for de novo chlamydial protein synthesis. End1 cells were infected with *CtrL2* at an MOI=5, treated with 50µg/ml chloramphenicol at 6hpi, and 24hpi levels of *TNF* transcripts and secreted TNF were monitored. As shown in Fig. 3A & B, both TNF expression at the transcript and secreted protein levels were enhanced by nearly 400- and 6-fold, respectively, in comparison to infected/mock-starved samples. These results indicate that enhanced TNF expression is activated directly or indirectly by a *de novo* synthesized chlamydial factor.

**Figure 3.**
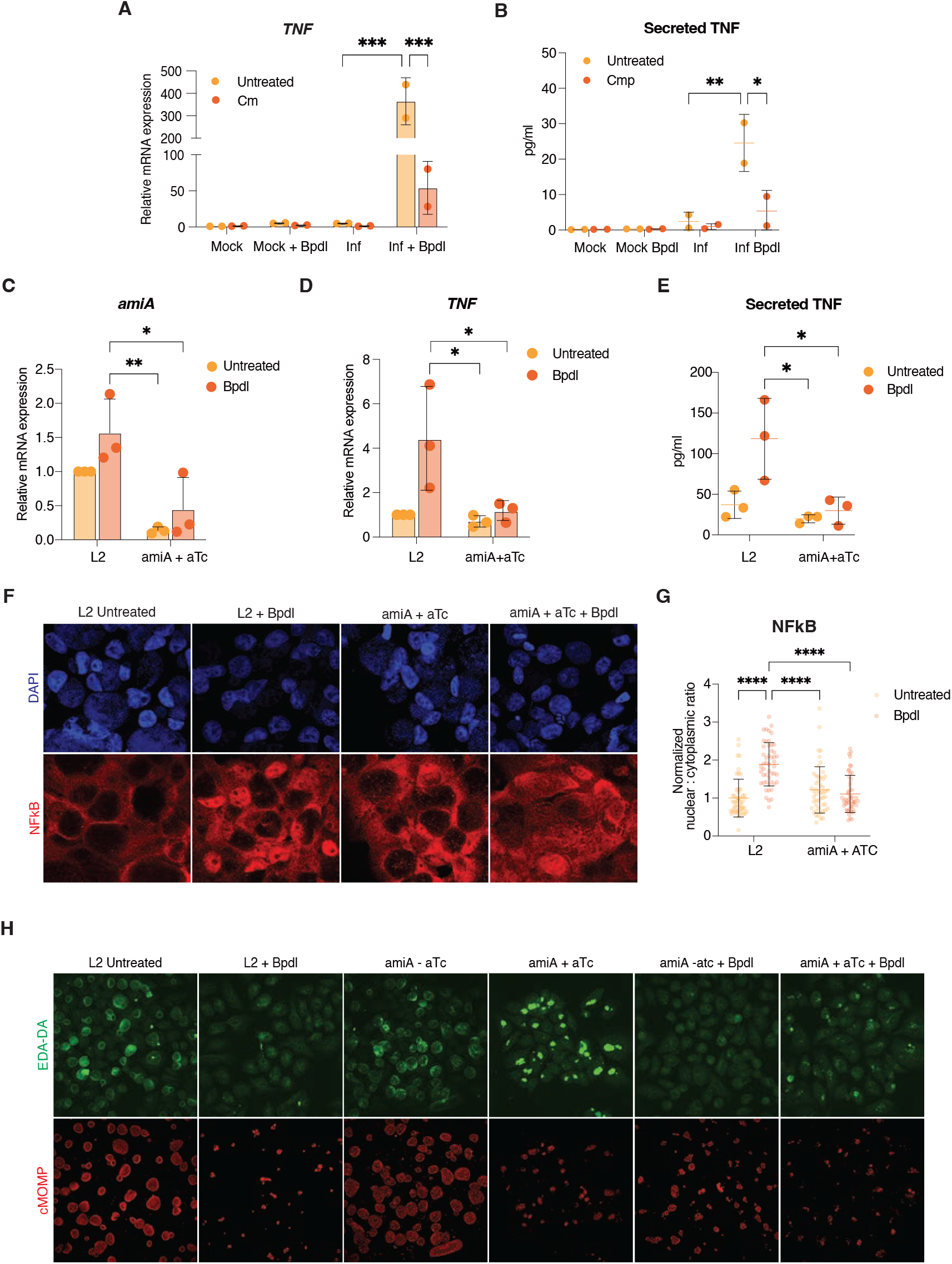
Inhibition of chlamydial proteins or peptidoglycan remodeling enzyme alters TNF expression in infected epithelial cells. **A)** *TNF* mRNA expression at 24hpi in End1 cells either mock infected or infected with *CtrL2*, treated with chloramphenicol at 6hpi and bipyridyl at 8hpi. Gene expression was quantified by qPCR (n=3). B) TNF protein levels secreted in the supernatants of mock-infected or *CtrL2*-infected End1 cells following treatment with chloramphenicol (Cm) and bipyridyl at 6hpi and 8hpi respectively, measured using Multiplexing LASER Bead Technology (n=3). **C)** Expression of *amiA* transcripts measured by qPCR in End1 cells infected with either *CtrL2* or *LCria amiA* strains, induced by anhydrous tetracycline at 8hpi (n=3). **D)** TNF mRNA levels at 24hpi measured by comparative qPCR in End1 cells infected with CtrL2 or *LCria amiA*, and either induced by anhydrous tetracycline or treated with bipyridyl at 8hpi (n=3). **E)** Quantification of TNF secreted in the supernatants of End1 cell at 24hpi, following infection with CtrL2 or LCRiaamiA and subsequent treatment with aTC or bipyridyl at 8hpi (n=3). **F)** Representative immunofluorescence micrographs showing NFκB (red) translocation to the nucleus (blue) in *CtrL2*- or *LCria amiA*- infected End1 cells treated with either bipyridyl or aTC at 8hpi and fixed at 24pi. **G)** Quantification of NFκB nuclear translocation in F) expressed as nuclear to cytoplasmic fluorescence intensity ratios. Data represent measurement from 50 randomly selected cells per treatment group. H) CtrL2-infected or *LCria amia*-infected (red) End1 cells were grown in the presence of EDA-DA (green) for 24h and treated with bipyridyl or aTC 8hpi. Images were selected from 5 fields across two-independent biological replicates.

Our previous characterization of the *CtrL2* transcriptome under iron-limiting conditions identified *amiA* as upregulated; and its role in PG remodeling raises the possibility of a role in NOD2 activation (24). In this regard, increased *amiA* transcription during Bpdl treatment was prevented by CRISPRi-mediated knockdown. In *Chlamydia* this system relies on constitutive expression of a guide RNA (gRNA) and inducible expression of a catalytically dead dCas9 by treatment with 10nM anhydrous tetracycline (aTC). The gRNA was designed to bind the promoter region of *amiA* for recognition and binding by dCas9, thus blocking the transcription machinery. Knockdown was monitored by RT-qPCR of endogenous transcripts and found to be efficient, with levels of reduction reaching 0.2-fold at 16h post-aTC induction (Fig. 3C). The effect of *amiA* knockdown on *TNF* transcription, TNF production/secretion, and NF-κB nuclear localization were simultaneously quantified in mock-starved and Bpdl-treated samples. Fig. 3C showed efficient knockdown upon aTC induction in mock-starved groups. An increase in *amiA* expression relative to mock-starved samples was observed in Bpdl-treated group. Inhibition of amiA upregulation during iron starvation was also observed. In parallel samples, *TNF* transcription, TNF secretion, and NF-κB nuclear translocation were decreased. Therefore, the basis for the pathogen driven NF-κB activation and enhanced TNF expression is the increased expression (i.e., transcriptional dysregulation) of *amiA*.

AmiA is an amidase that is involved in PG degradation and remodeling (38). Specifically, it cleaves the tetrapeptide motif that includes the terminal D-Ala-D-Ala dipeptide. Using a click chemistry-compatible EDA-DA analog, the *CtrL2* PG network was fluorescently labeled, resulting in highly localized staining profiles at steady state conditions that reflect the presence of PG only in the division septum (39). Of note, pathogenic chlamydiae lack a PG sacculus. We speculated that dysregulated *amiA* overexpression in iron-starved *CtrL2* would result in increased AmiA activity, with the subsequent cleavage of EDA-DA resulting in changes to label localization. Indeed, as shown in Fig. 3H, *CtrL2*-localized EDA-DA signal was lost in Bpdl-treated samples. In control mock starved groups, transformants harboring the amiA CRISPRi plasmid showed similar localization profiles as the parental *CtrL2* strain. Induction of knockdown (AmiA+aTC) maintained localization within the bacteria, albeit at reduced levels. The reduction in the overall fluorescence signal could be attributed to effects on bacterial cell division, as reflected by the smaller inclusions. Noteworthy was the effect of inhibiting *amiA* induction in Bpdl-treated *CtrL2*, where retention of EDA-DA signals within the bacteria was observed. Taken together, the data indicate that *amiA* dysregulation during iron starvation leads to the loss of retention of D-Ala-D-Ala PG intermediates within bacteria, NFκB activation, which depends on NOD2, and enhanced *TNF* transcription and cytokine secretion.

Recalling observations that *CtrL2*-infected End1 cells upon iron starvation exhibited increased cell death, we investigated whether inhibiting *de novo* protein synthesis would affect LDH levels – the proxy for cell death, in the growth media. As shown in Fig. 4A, inhibition of chlamydial protein synthesis led to a significant decrease in LDH level in infected iron-starved samples. Furthermore, suppression of NOD2 signaling pathway in *CtrL2*-infected, iron-starved cells also resulted in reduced LDH levels (Fig. 4B). These findings suggest that increased cell death in iron-starved and *CtrL2*-infected epithelial monolayers is linked to dysregulated expression of PG remodeling enzymes, the subsequent activation of the NOD2 signaling pathway, and enhanced NF-κB-dependent TNF expression.

**Figure 4.**
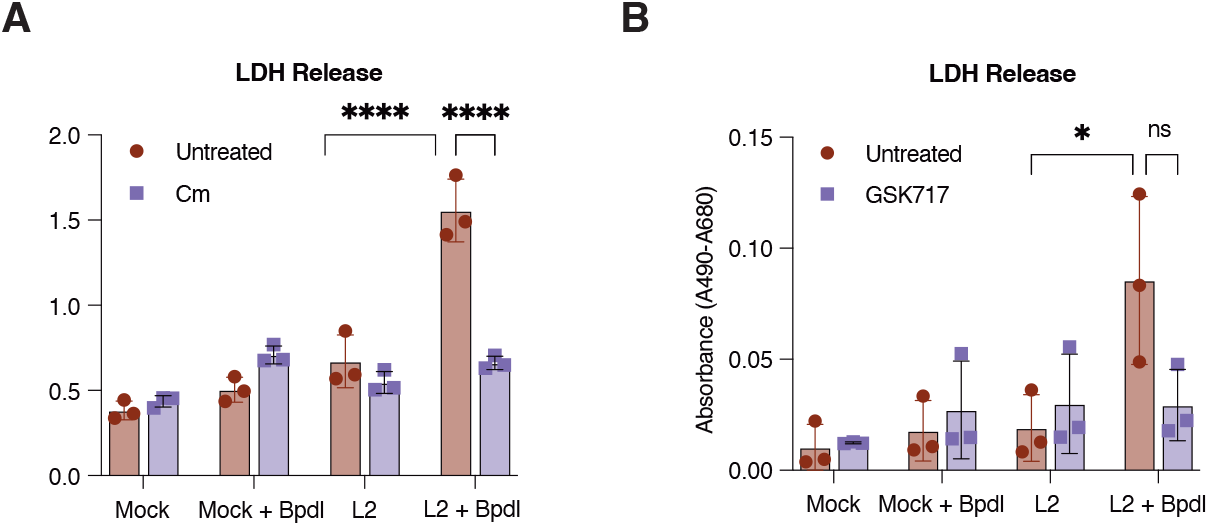
Inhibition of chlamydial protein synthesis and NOD2 signaling pathway reduces LDH release under *CtrL2-i*nfected iron-starved conditions. **A)** LDH release measured by absorbance in chloramphenicol treated End1 cells either mock-infected or infected with *CtrL2*, with or without bipyridyl treatment at 24hpi (n=3). **B)** LDH release as measured by absorbance in supernatants from GSK717-pretreated End1 cells infected with *CtrL2* and subjected to 16h of iron starvation at 24hpi (n=3). Statistical significance determined by two-way ANOVA and Tukey’s multiple comparisons test. ****p<0.0001; *p<0.03; ns, not significant.

## Discussion

Here, we provide evidence that nutritional immunity in the form of iron starvation stimulates pathogen recognition and consequent death of infected cells, thus providing an additional mechanism of how nutrient deprivation contributes to the elimination of the pathogen. While starving the pathogen of iron contributes to pathogen elimination, we provide a more refined model of nutritional immunity by implicating iron starvation in gene expression “errors” in the pathogen to activate pathogen recognition signaling in infected cells.

Chlamydiae reside within cells in pathogen-modified vacuoles, called inclusions, with the inclusion membrane serving as the host-pathogen interface for communication and subcellular interactions with the host cell. Subcellular components that interact with the chlamydial inclusion include nutrient-laden vesicles, organelles, and signaling proteins (40). With the membrane’s limited permeability to molecules that are >700 daltons, it also serves as a sieve to regulate entry and exit of molecules. For example, the limited permeability of the inclusion membrane likely plays a role in chlamydial iron homeostasis, with iron atoms transported to the inclusion via the slow pathway of transferrin receptor recycling being inaccessible to the host high-affinity iron-sequestering ferritins (41). This is crucial given that *C. trachomatis* is incapable of synthesizing high-affinity siderophores that can compete for iron binding with host cell ferritins. This limited two-way permeability of the inclusion membrane could additionally restrict exit of PAMPs, with molecular weight above 700 daltons (42).

PG recycling in *C. trachomatis* remains poorly understood with only characterized function attributed to NlpC/60, which hydrolyzes PG-derived peptides and recycles PG ring turnover products (43). In model prokaryotes, such as *E. coli*, anhydrodisaccharides generated by amidase action on anhydrous muropeptides, are recycled from the extracellular environment to the bacterial cytosol by AmpG and/or YejBEF complexes, where they are fed into the biosynthetic pathway (44). The tetrapeptide product is captured by MppA for transport into the bacterial cytosol by the OppBCDF complex (45, 46). *C. trachomatis* has homologs of AmpG, YejBEF, and the OppBCDF complexes (47). We envision that an overactive AmiA leads to the overaccumulation of anhydrous muropeptides, thus overwhelming the recycling machinery, and triggering NOD2 recognition of PAMPs. While reduction of expression of one or more components of the recycling machinery essentially could also result in PAMP accumulation, changes to their expression were not observed in the transcriptome of Bpdl-treated *C. trachomatis*. These observations are consistent with AmiA overexpression as a major contributor to the activation of NOD2/ NF-κB/TNF axis, as indicated by *amiA* knockdown being sufficient to eliminate signaling activation.

A particularly interesting aspect is the sufficiency of NOD2 inhibition by GSK717 in neutralizing NF-κB activation and overexpression of TNF. Our findings are consistent with a previous report on the absence of NOD1 activation in Bpdl-treated *C. trachomatis* (48). Liechti *et al*. did not address NOD2 signaling. Our data indicate that NOD2 recognition of *C. trachomatis* may be relevant in iron-starved conditions. Specific inhibition of NOD2 by GSK717 or knockdown of AmiA was sufficient to reduce NF-κB nuclear translocation, and therefore no meaningful contributions from other pathways, either from the host or pathogen, were necessary. Collectively, our data point to a very exclusive association between iron starvation and pathogen recognition by NOD2. This association with a very specific type of nutrient starvation could potentially explain the paucity of reports implicating NOD2 in chlamydial detection by epithelial cells. If detection by NOD2 applies only to a specific growth condition, (*i*.*e*., iron starvation), then what might be the implications regarding acquisition of anti-NOD2 mechanisms? Such a specialized mechanism is unlikely to have been prioritized through evolution, thus identifying a potential therapeutic strategy that enhances NOD2-dependent recognition of *C. trachomatis*. An unexpected observation was the absence of activation of the NOD2/ NF-κB/TNF signaling axis in *C. trachomatis* strains overexpressing AmiA from a plasmid to bypass the need for Bpdl treatment. This discrepancy could be explained by the additional effects of iron starvation on both pathogen and host epithelial cell. With the latter, we observed a 17-fold increase in NOD2 expression (24), but not NOD1, TLR2, TLR1, etc., thus, potentially sensitizing the infected cell to NOD2 activation. Inappropriate production of other NOD2-recognized PAMPs by iron-starved chlamydiae, as a consequence of amiA dysregulation, could also contribute to activation of NOD2 and downstream signaling components. PG synthesis is highly regulated during chlamydial cell division, with the necessary PG biosynthetic, remodeling, and recycling genes limited during cell division. Dysregulation of one or more of these sets of genes arising from iron starvation could lead to enhanced pathogen recognition. While temporal restriction and recycling could also facilitate evasion of pathogen recognition by the host cell, evidence obtained supports amiA transcriptional regulation as the important rate-limiting process in evading recognition by NOD2.

Our findings raise the question of biological relevance of iron starvation in *C. trachomatis* during *in vivo* infection. One scenario is the lactoferrin expression, which fluctuates during female menstruation (49, 50). Lactoferrin present in the mucosa sequesters iron; and various mucosal pathogens, such as *Neisseria gonorrhoeae*, rely on lactoferrin receptors on their surface to obtain iron from this pool (51). There also have been correlative studies that assigned an inhibitory role for lactoferrin in *C. trachomatis* infections (52, 53). In addition, susceptibility to *C. trachomatis* infection is known to correlate with specific stages of the menstrual cycle (54, 55). Therefore, dysregulated *amiA* expression when high levels of lactoferrin are present could occur, with enhanced pathogen recognition manifesting as decreased infection. It is tempting to speculate that alterations of the cytokine profile could have significant downstream consequences on neighboring cells, resident innate immune cells, and the overall nature of the immediate and long-term immune response of the host.

In summary, the consequence of nutritional immunity extends beyond killing of pathogens by starvation. Instead, it is more nuanced with dysregulation of pathogen gene expression at multiple levels compromising its ability to deploy its survival strategies, including immune evasion.

## Materials and Methods

### Cell culture of cervical epithelial cells

Dulbecco’s Modified Eagle Medium (DMEM; Gibco, Thermo Fisher Scientific) supplemented with 10% (vol/vol) fetal bovine serum (Tissue Culture Biologicals), 2mM L-glutamine and 10µg/ml gentamycin was used to culture human female cervical epithelial adenocarcinoma HeLa cells that were obtained from ATCC (CCL-2.1). Additionally, keratinocyte serum-free medium (KSFM) (Thermo Fisher Scientific 37010022) supplemented with bovine pituitary extract, human EGF and, 0.4mM CaCl2 was used to maintain human endocervical epithelial HPV-16 E6/E7-immortalized End1s (End1 E6/E7, ATCC CRL-2615). All experiments were performed using cells at passage numbers between 3 and15.

### *Chlamydia* Infection, strains and treatment conditions

*Chlamydia trachomatis* serovar L2 (434/Bu), originally obtained from Dr. Ted Hackstadt (Rocky Mountain Laboratories), was propagated in McCoy cells, and elementary bodies (EBs) were purified via density gradient ultracentrifugation as previously described (56). HeLa cells at approximately 80% confluency were washed with HBSS (Thermo Fisher Scientific, 14025134) and infected using an inoculum prepared in unsupplemented DMEM at the desired multiplicity of infection (MOI). To synchronize infection, cells were centrifuged at 500 × g for 15 minutes at 4°C using an Eppendorf 5810 R tabletop centrifuge with an A-4-81 rotor. Following centrifugation, the inoculum was removed and replaced with fresh, pre-warmed complete medium, and cells were returned to the incubator until the indicated time points. To induce iron starvation, the culture medium was replaced at 8 hours post-infection (hpi) with medium containing 100 µM bipyridyl, and cells were maintained under these conditions for 16 hours. Inhibition of Chlamydia protein synthesis was achieved by adding chloramphenicol at a final concentration of 50 µg/mL at 6 hpi. For inhibiting NOD2 signaling, cells were pretreated with 15 µM GSK717 (Millipore Sigma, 5337180001) for 2 hours prior to infection, and the compound was maintained in the medium for the duration of the experiment. *C. trachomatis LCria amiA*, kindly provided by Dr. Scot Ouellette (University of Nebraska Medical Center), were infected at an MOI of 1 following the same protocol described above. Where indicated, anhydrotetracycline was added at a final concentration of 10nM at 8hpi. For infections in End1 cells, cultures were initiated at ∼125% confluency and infected using the same protocol.

### RNA isolation, cDNA prep and quantitative PCR

RNA was extracted from *L2*-infected or uninfected cells treated with or without Bipyrydyl. Cells were scraped from a 6-well plate using 500µL of TRIzol reagent (Thermo Fisher Scientific 15596026.) pre-chilled on ice. The resulting lysates were transferred into RNase-free tubes containing approximately 100µL of zirconia beads to disrupt cells and inclusions. Samples were vortexed for 10 minutes, then centrifuged at 21,000 × g for 10 minutes at 4 °C to pellet the beads. The supernatant was transferred to a fresh tube containing 100 µL of chloroform, vortexed, and incubated at room temperature for 10 minutes. Following a second centrifugation at 21,000 × g for 15 minutes at 4 °C, the aqueous phase was carefully collected and mixed with 250 µL of ethanol. RNA was then purified using the PureLink RNA Mini Kit (Thermo Fisher Scientific, 12183018A) and cDNA was synthesized using SuperScript IV VILO (Thermo Fisher Scientific, 11756050). The resulting cDNA was diluted 1:10 for quantitative PCR (qPCR) reactions. qPCR was performed using TaqMan Universal master mix (Thermo Fisher Scientific 4440038) to analyze host eukaryotic genes, with TaqMan probes (Thermo Fisher Scientific 4331182) and HPRT1 (Hs02800695_m1.) as the housekeeping gene. A list of target genes is provided in Table S1. For the analysis of *Chlamydia trachomatis* genes, SYBR Green Master Mix (Thermo Fisher Scientific 4364344.) was used, and expression levels were normalized to either *groEL* or *euo*. Primer sets are included in Table S1. Relative gene expression as fold changes were calculated using the comparative Ct (ΔΔCt) method as described (57).

### siRNA transfection of NF_κ_B

Transfections were carried out in HeLa cells, which exhibit greater permissivity to siRNA compared to End1 cells. HeLa cells at approximately 50% confluency were transfected with SignalSilence Control siRNA (Cell Signaling Technology 6201S) or SignalSilence NF-κB p65 siRNA (Cell Signaling Technology 6261S) using RNAiMAX (Thermo Fisher Scientific, 13778030), following the manufacturer’s protocol, at an optimized concentration of 10nM. At 24h post-transfection cells were infected with *L2* at an MOI of 5 followed by treatment with bipyridyl at 8hpi. At 24hpi, cells were lysed for RNA extraction or western blot analysis. siRNA-mediated knockdown was confirmed by both qPCR at the transcript level and western blotting at the protein level. All experiments were conducted in 6-well plates.

### Lysate preparation and Immunoblotting

HeLa or End1 cells cultured in 6-well plates were infected with *L2* and treated with bipyridyl. At 24hpi, cell lysates were collected using hot 1% SDS lysis buffer supplemented with protease and phosphatase inhibitors to prevent post-CPAF-mediated effects, as described in (58). The lysates were treated with Pierce Universal Nuclease (1:1000 dilution, 88701) for 5 minutes at room temperature to degrade nucleic acids. Subsequently, 4X Laemmli Sample Buffer containing β-mercaptoethanol (1:10 dilution) was added. Samples were boiled at 95°C for 5 minutes before being loaded onto a 12% acrylamide/bis-acrylamide gel and transferred to a nitrocellulose membrane (Bio-Rad) using a semi-dry transfer method. Membranes were blocked with 5% non-fat dry milk for 1 hour, then incubated overnight at 4°C with primary antibodies: NF-κB p65 (D14E12) XP® (Cell Signaling Technology, 1:1000), Anti-TNF alpha antibody [EPR19147] (Abcam, 1:1000, ab183218), β-Actin (Cell Signaling Technology, 3700S, 1:5000), Vimentin (, followed by 2-hour incubation at room temperature with IR dye-conjugated secondary antibodies: Donkey Anti-Rabbit IgG Polyclonal Antibody IRDye® 800CW (LI-COR Biosciences, 926-32213, 1:10000) and Donkey Anti-Mouse IgG Polyclonal Antibody IRDye® 680RD (LI-COR Biosciences, 926-68072 1:10000). Blots were visualized using the Azure Biosystems c600 under near-infrared (NIR) settings. Images were analyzed, processed, and quantified using ImageJ.

### Cytokine Profiling

Supernatants from each well of the 6-well plates were collected and transferred to 2 mL Eppendorf tubes, then centrifuged at 21,000 × g for 5 minutes at 4°C to pellet any floating dead cells. A volume of 350 µL from each supernatant was sent to Eve Technologies Pvt. Ltd. (Canada) for cytokine analysis using a multiplex flow cytometry-based assay. Human Cytokine/Chemokine 96-Plex Discovery Assay Array® (HD96) was used for this purpose.

### Immunofluorescence Assay (IFA)

HeLa cells were either mock-infected or infected with *Chlamydia trachomatis L2* on acid-washed coverslips placed in a 24-well plate. Where indicated, cells were treated with bipyrydyl, chloramphenicol, GSK717, or anhydrous tetracycline. At the designated timepoints, cells were fixed with 4% paraformaldehyde and permeabilized using 0.2% Triton X-100 (Cat. No.). Following fixation and permeabilization, cells were blocked with 3% BSA in PBST for 1 hour and incubated overnight at 4°C with primary antibodies: NF-κB p65 (D14E12) XP® (Cell Signaling Technology, 1:500), MOMP (Meridian Life Sciences, B65266G, 1:1000) and cHSP60 (Thermo Fisher Scientific MA3-023, 1:1000) wherever indicated followed by incubation with secondary antibodies. Between each step, cells were washed at least three times with PBST. Nuclear staining was performed using 2.5 μg/mL DAPI for 5 minutes, followed by a PBST wash. Coverslips were then mounted onto slides using ImmunoMount and allowed to dry undisturbed overnight. Images were acquired using a Nikon Ti2 Eclipse spinning-disk confocal microscope and subsequently processed and quantified using ImageJ.

### Quantification of Nuclear to Cytoplasmic ratio of NF_κ_B using Image J

Images were acquired using a Nikon CSU-W1 spinning disk confocal microscope, capturing a minimum of five fields per coverslip. Ten nuclei were randomly selected based on DAPI staining, and their outlines were manually masked. Corresponding cytoplasmic regions were also manually masked using the NF-κB channel. Nuclear-to-cytoplasmic intensity ratios were calculated and normalized to the mean ratio of mock-infected, untreated controls for relative comparison. Quantification was performed across three independent experiments using ImageJ.

### EDA-DA staining of the peptidoglycan

Peptidoglycan staining was performed using a click chemistry reaction as previously described (39). Briefly, EDA-DA (Tocris, 7714) was added at a final concentration of 0.5 mM at the time of infection and maintained throughout the duration of the experiment. At specified timepoints, appropriate treatments were applied, and cells were fixed with methanol and permeabilized using 0.2% Triton X-100. Blocking was carried out with 3% BSA for one hour. The Click-iT kit (Thermo Fisher Scientific, C10269) was used according to the manufacturer’s instructions, followed by an additional 30-minute wash with 3% BSA. Cells were then stained for MOMP and DAPI to detect *C. trachomatis* and cell nuclei, respectively, using the procedure outlined above. Imaging was performed on a Nikon Ti2 Eclipse spinning-disk confocal microscope, and images were processed using ImageJ.

### Cell death Assay

Supernatants from both mock-infected and *L2*-infected samples were combined in 2 mL Eppendorf tubes and centrifuged at 21,000 × g for 15 minutes at 4 °C to pellet dead cells. Cytotoxicity was assessed using the CyQUANT™ LDH Cytotoxicity Assay Kit (Thermo Fisher Scientific, C20301) according to the manufacturer’s instructions, with absorbance measured at 490 nm and 680 nm. For the live/dead assay involving Annexin V staining, the Apoptotic, Necrotic, and Healthy Cells Quantitation Kit (Biotium, 30066) was used following manufacturer’s protocol. Cells were seeded in 24-well plates, and after staining, imaged using a Zeiss fluorescence microscope with a 20X objective. Images were processed using Image J.

### Graphs and Statistical analysis

All graphs and statistical analyses were conducted using GraphPad Prism version 10.4.2 (GraphPad Software, Inc.). Experiments were performed in three independent biological replicates, and data are presented as mean ± standard deviation (SD). Statistical significance was assessed using one-way ANOVA for comparisons involving more than two groups and two-way ANOVA for grouped data, followed by Tukey’s post hoc test. A p-value of less than 0.05 was considered statistically significant.

## Acknowledgments

We would like to acknowledge all the members of Carabeo lab for their valuable feedback on the analysis and design of the experiments.

---

**Fig. S1.**
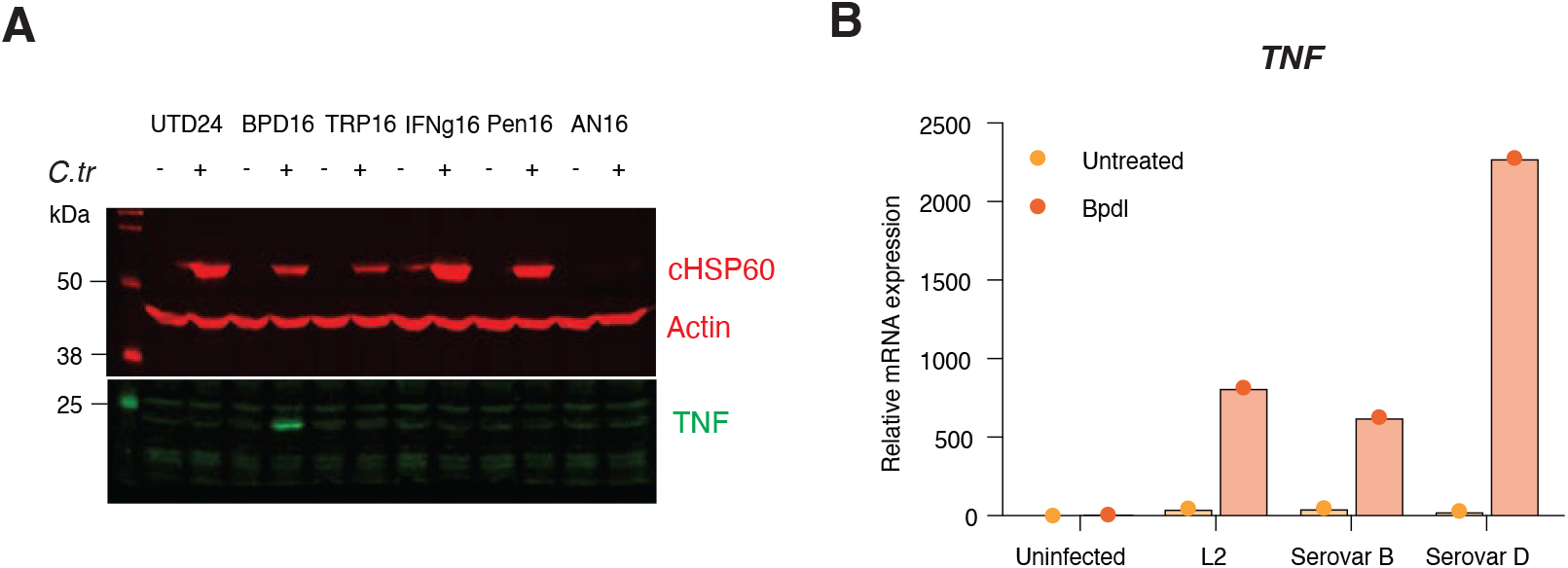
Enhanced TNF expression observed in iron starved infected epithelial cells. **A)** Western blot analysis of intracellular TNF at 24hpi in HeLa cells either mock or infected with *CtrL2* treated at 8hpi with several stressors i.e. Bipyridyl (100uM), tryptophan starvation (DMEM w/o tryptophan), IFN-g treatment (200U/ml), Penicillin G (0.6ug/ml) and AN3365 leucyl-tRNA synthetase inhibitor (5ug/ml) (n=1). **B)** TNF mRNA expression in HeLa cells infected with *CtrL2, CtrB* and *CtrD* at 24hpi analyzed by RT-qPCR normalized to HPRT1 under iron starved conditions induced 8hpi using bipyridyl (n=1).

**Fig. S2.**
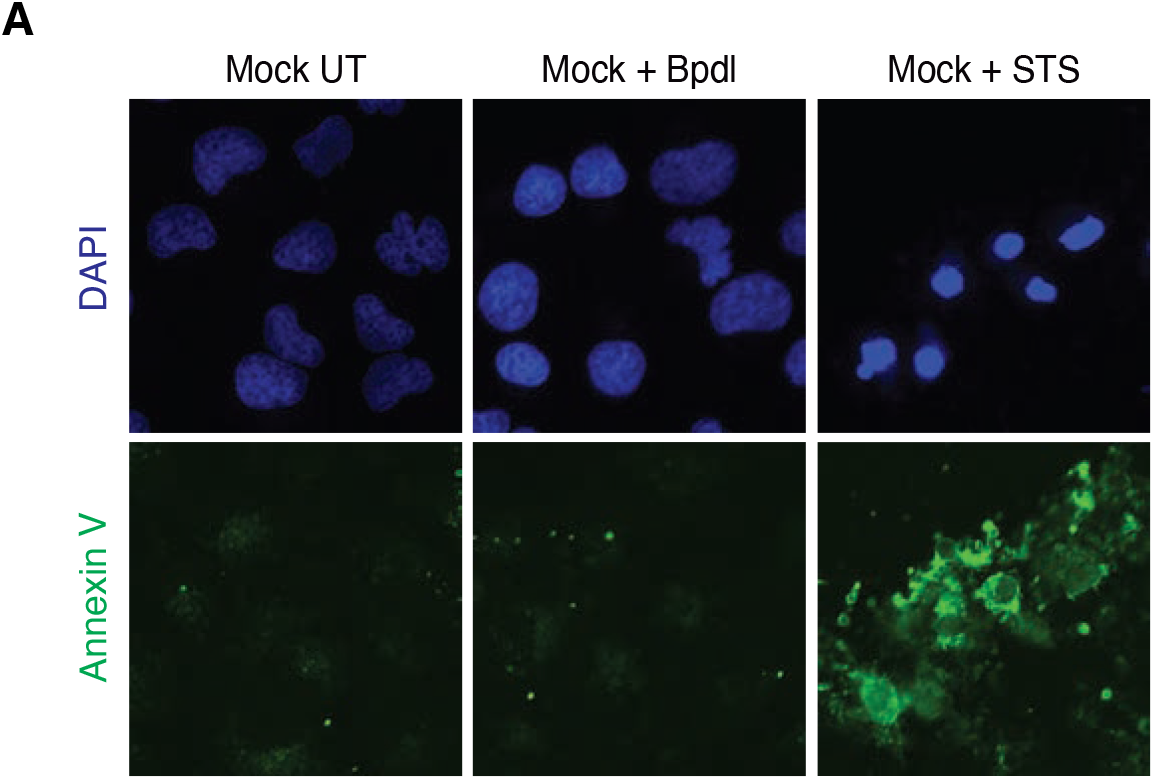
Annexin V staining in Mock-infected HeLa cells. **A)** Representative micrographs of HeLa cells stained with Annexin V at 24h following mock infection and bipyridyl treatment (n=2).

**Supplementary Table 1.**
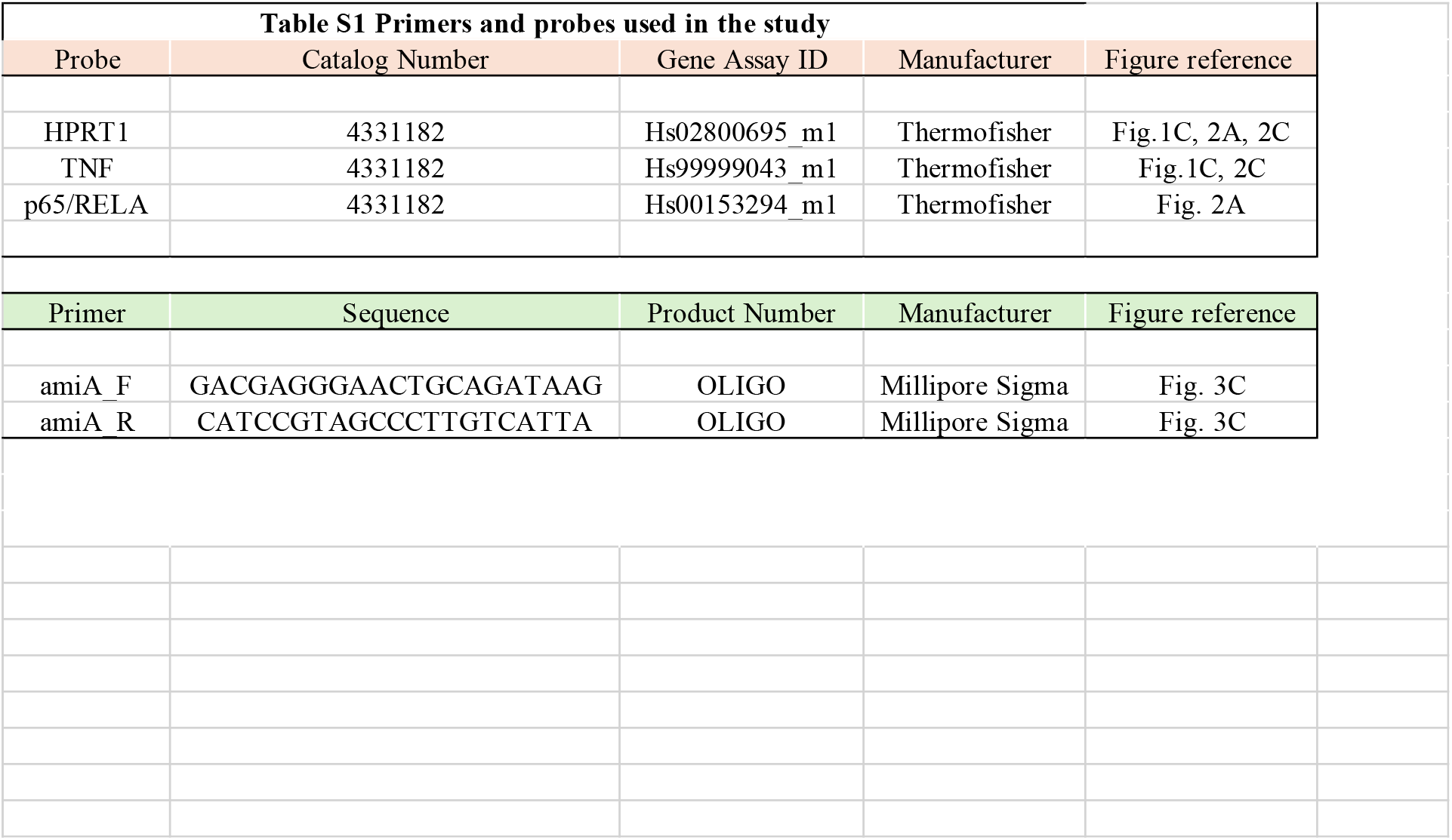
List of probes and primers used in the study.

